# Whole brain mapping of somatosensory responses in awake marmosets investigated with ultra-high field fMRI

**DOI:** 10.1101/2020.08.05.238592

**Authors:** Justine C. Cléry, Yuki Hori, David J. Schaeffer, Joseph S. Gati, J. Andrew Pruszynski, Stefan Everling

**Author notes:** **Corresponding authors:** Stefan Everling, PhD, Centre for Functional and Metabolic Mapping, Robarts Research Institute, 1151 Richmond Street North, London, Ontario, N6A 5B7, Canada., Justine C. Cléry, PhD, Robarts Research Institute, 1151 Richmond Street North, London, Ontario, N6A 5B7, Canada. **Author contribution:** J.C.C., J. A.P. and S.E. designed research; J.C.C., Y.H., D.J.S, J.S.G. and S.E. performed research; J.C.C. analyzed data, prepared the figures and wrote the paper; and J.C.C., Y.H., R.S.M., J.A.P. and S.E. edited the paper.

## Abstract

The common marmoset *(Callithrix jacchus)* is a small-bodied New World primate that is becoming an important model to study brain functions. Despite several studies exploring the somatosensory system of marmosets, all results have come from anesthetized animals using invasive techniques and post-mortem analyses. Here we demonstrate the feasibility for getting high-quality and reproduceable sensorimotor mapping in awake marmosets with functional magnetic resonance imaging (fMRI). We acquired fMRI sequences in four animals while they received tactile stimulation (via air-puffs), delivered to the face, arm or leg. We found that the body representation progressed medially from the leg to the face in areas 3a, 3b, 1/2, and from caudal to rostral sites in areas S2 and PV. SI and SII exhibited a body representation in their functional connectivity pattern within the posterior and midcingulate and the thalamus. Interestingly, we also found a somatotopic body representation in two subcortical areas: the thalamus and, for the first time, in the putamen. These maps have similar organizations as those previously found in Old World macaque monkeys and humans, suggesting that these subcortical somatotopic organizations were already established before Old and New World primates diverged. Our results show the first whole brain mapping of somatosensory responses acquired in a non-invasive way in awake marmosets.

**Significant statement:** Here we used somatosensory stimulation combined with functional magnetic resonance imaging to map whole brain activation in awake marmosets. We used light tactile stimulation, consisting of air-puffs, delivered on the face, arm or leg. We found a topographic body representation in primary (SI) and secondary (SII) somatosensory regions, thalamus and putamen. We also revealed the existence of a body representation organization within the thalamus and the cingulate cortex by computing functional connectivity maps from seeds defined in SI/SII for face, arm and leg using resting-state fMRI data. This non-invasive approach will be essential for chronic studies by guiding invasive recording and manipulation techniques.

## Introduction

The somatosensory system and tactile processing have been studied for several decades in numerous mammals (Kaas, 1987, 1989, 2004a, 2004b; Krubitzer, 1996). Two main cortical regions have been identified in the somatosensory system, the primary somatosensory area (SI) and the secondary somatosensory area (SII). The number of subdivisions in these regions varies between mammalians species (Kaas, 1987, 1989). In humans and non-human primates, unlike rodents (for review, see Kaas, 2004a), SI is subdivided into 4 architectonic fields: areas 3a, 3b, 1 and 2, which contain a separate but parallel topographic body representation (Kaas and Pons 1988). SII contains 4 subdivisions but has only a single body representation (Kaas et al., 1979).

The common marmoset *(Callithrix Jacchus),* a New-World primate, has received much attention in neuroscientific research field in recent years (Burkart and Finkenwirth, 2015; Silva, 2017; Preuss, 2019). This is primarily because of: 1) ease of breeding in captivity and manageability as experimental animals (small body size, less biosafety concerns) (Mansfield, 2003); 2) long lifespan useful in the research of development and aging (Takahashi et al., 2015; Workman et al., 2019); 3) feasible transgenic techniques (Sasaki et al., 2009), and 4) a lissencephalic brain, amenable to electrophysiological recording owing to easy access to target area. In the context of the somatosensory system, several previous studies have established that, like humans and other non-human primates, both areas SI and SII in marmosets have a somatotopic body representation (Carlson et al., 1986; Krubitzer and Kaas, 1988, 1990, 1992; Garraghty et al., 1990; Huffman and Krubitzer, 2001a, 2001b; Qi et al., 2002; for review, see Kaas, 2004b). To date, somatotopic studies in marmosets have been performed under anesthetized conditions and/or using post-mortem analyses. One exception is the recent studies performed by Silva and his colleagues who have demonstrated functional magnetic resonance imaging technique (fMRI) in conscious, awake marmosets using habituation procedures (Silva et al., 2011) and explored the differences of brain activation in response to somatosensory stimulation between awake and anesthetized conditions (Liu et al., 2013; Papoti et al., 2013; Hirano et al., 2018). Although these researches were focused on only responses evoked by electrical stimulation on the wrist, they observed significantly attenuated responses in SI, SII, and thalamus or abolished responses in caudate nucleus under anesthetized compared to awake conditions. These results suggest that the somatotopic maps obtained in anesthetized monkeys may underrepresent the reality of the tactile processing occurring at an awake state. As such, it is important to measure the somatotopic representation in awake marmosets.

In addition, connectivity analyses in marmosets have been mainly limited to post-mortem analyses using anatomical tracers (Krubitzer and Kaas, 1990, 1992; Huffman and Krubitzer, 2001a, 2001b; Qi et al., 2002). A recent study showed that functional connectivity (FC) from resting-state fMRI (RS-fMRI) data was highly correlated with tracer based cellular connectivity data, even though some variabilities were observed across seed regions (Hori et al., 2020a). FC from RS-fMRI seems to be appropriate to explore the tactile connectivity mapping in awake marmosets to understand how somatosensory information is relayed within the brain.

Here we performed whole brain mapping of somatosensory responses in the awake common marmoset by using high resolution fMRI in a 9.4T combined with tactile stimulation evoked by air puffs. The purpose of this study was threefold. First, to establish the feasibility for getting high-quality and reproducible sensorimotor mapping with fMRI acquired in awake condition. Second, to validate the existence of the somatotopic body representation patterns observed in anesthetized animals, by applying tactile stimulation to the face, arm or leg of awake marmosets. Third, to explore how somatosensory areas highly responsive to the tactile stimulation are functionally connected to other cortical and subcortical regions.

## Methods

All experimental methods described were performed in accordance with the guidelines of the Canadian Council on Animal Care policy on the care and use of experimental animals and an ethics protocol approved by the Animal Care Committee of the University of Western Ontario. Monkeys were under the close supervision by the university veterinarians.

### 1. Somatosensory system through task-based fMRI study

#### 1.1. Subjects and experimental setup

Four common marmosets *(Callithrix jacchus)* were the subjects in this study (3 males, 1 female). Their ages and weights at the time of the experiments were 30 months and 390 g (M1), 26 months and 330 g, (M2) 28 months and 295 g (M3) and 24 months and 355 g (M4), respectively. The animals were prepared for awake fMRI experiments by implanting an MRI-compatible head restraint/recording chamber (for details, see Johnston et al., 2018). Each animal was initially habituated to the MRI environment over the course of three weeks. First, the marmosets were acclimatized to the animal holder (tube closed by a neck and tail plates), the head-fixation system, and the MRI sequence (for details, see Schaeffer et al., 2019). The behavior of the animals was monitored through the training to assess the tolerance and the well-being of the marmosets (Silva et al., 2011).

For the MRI acquisition, the animal’s left arm and left leg were shaved to apply more easily the tactile stimulation directly to the animal’s skin. The plastic contacts by where the tactile stimulus was delivered (stimulus node) were taped around the shaved arm and leg of the animal. The animal was placed in the sphinx position in an animal holder consisting of an MRI-compatible restraint system (Schaeffer et al., 2019) with a plastic manipulator arm at the front for delivering tactile stimuli to the face. The animal was first restrained using neck and tail plates, then the head was restrained by the fixation of the head chamber to the five-channel receive coil. An MRI-compatible camera (Model 12M-i, 60-Hz sampling rate, MRC Systems GmbH, Heidelberg, Germany) was positioned in front of the animal holder to monitor the animals during the acquisition sessions by a veterinary technician but was too far from the eyes to measure the eye movements or pupil dilatation. The tactile task onset was launched in synchronization with the first MRI trigger via a custom-written program running on a Raspberry Pi (Model 3 B+, Raspberry Pi Foundation, Cambridge, UK). All the MRI trigger and the tactile stimulation delivery outputs were recorded through LabChart 7 software (ADInstruments, Sydney, Australia).

#### 1.2. Tactile Stimulation

The tactile stimulation consisted of repeated air puffs delivered to the face (lower part of the cheek, encompassing parts of the chin, nose and mouth, Fig. 1A, left draw), to the upper body part (the exterior part of the forearm for M1, M2, M4, see Fig. 1A, middle draw, or the interior part of the forearm for M3) or to the lower body part (the back part of the leg, corresponding to the calf, Fig. 1A, right draw). Air puffs were delivered by the pneumatic Galileo Somatosensory™ stimulus system (Galileo Rack Mount Model, Epic Medical Concepts & Innovations, Inc., Mission, Kansas USA) using stimulus nodes wrapped around the arm/leg of the animal with veterinarian wrap, or via a manipulator arm in the front of the animal’s face (Fig. 1A). Only the left side of the animal was stimulated. The MRI-compatible stimulus nodes and flexible tubing were made of urethane and silicon, respectively. The internal diameter of the stimulus node was 6mm. The length of flexible tubing was 25’. The air pressure was set to 0.7 Bar.

**Figure 1:**
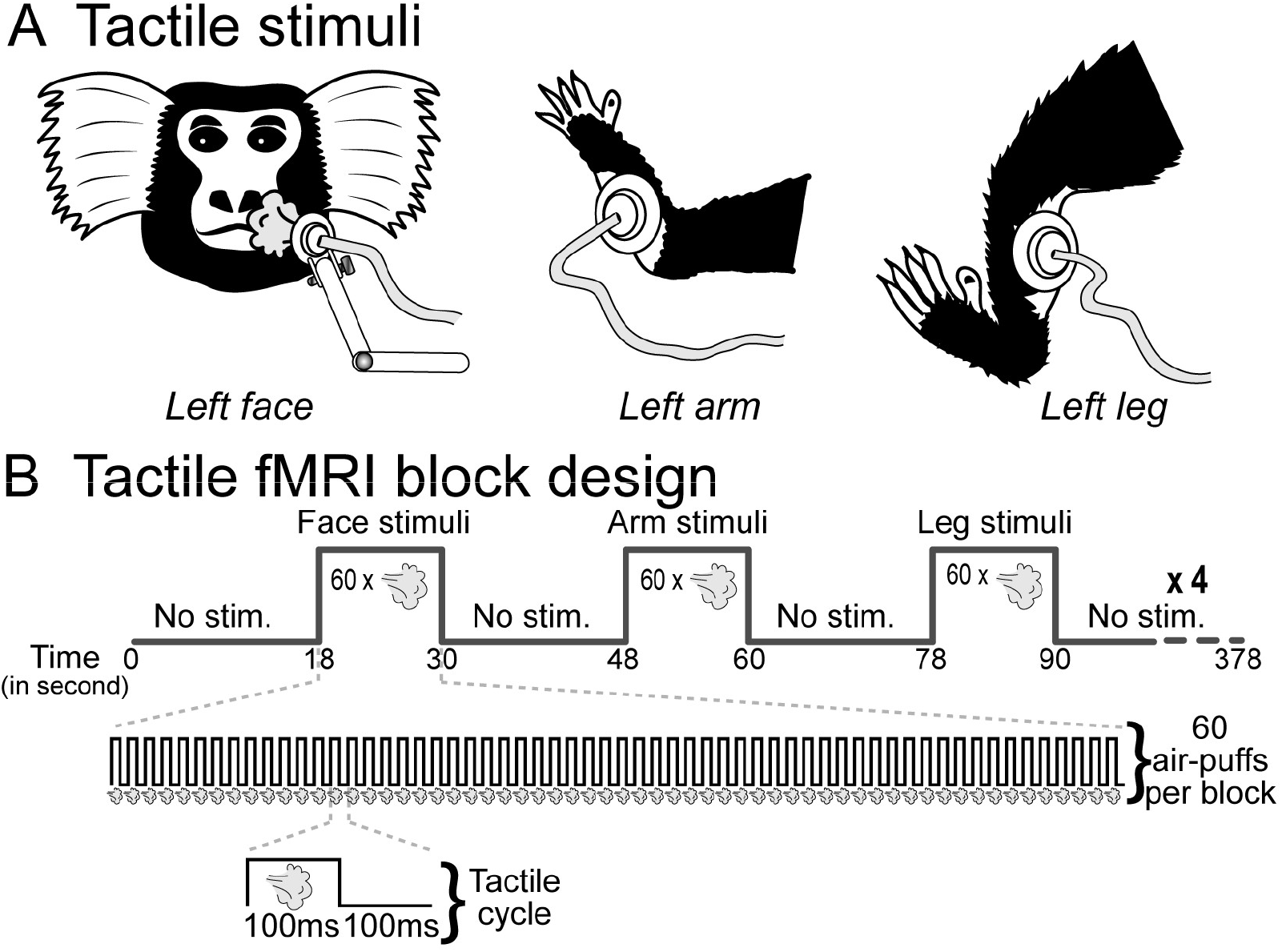
Experimental fMRI protocol. A) Tactile stimulation consisted of air-puffs delivered to the left face, arm or leg of the animals. B) Tactile fMRI block design. It consisted of a succession of blocks in which the face, the arm or the leg were stimulated in separate blocks (60 air-puffs delivered for a duration of 12 seconds per block). Blocks without any stimulation separated tactile blocks for 18 seconds. This pattern was repeated four times.

The task was built and ran with the Galileo software and tactile stimuli were delivered following a task-block design (Fig. 1.B). Each tactile stimulation block lasted 12s (8 functional volumes) and was separated by no stimulation blocks of 18s (12 functional volumes). The different tactile stimuli (face, arm and leg) were presented within the same run but in separate blocks. Each tactile stimulus lasted 100ms, was delivered every 200ms and repeated 60 times within a tactile stimulation block (Fig. 1B). Each type of tactile stimulation was repeated 4 times during the sequence (for a total duration of 6min 18sec or 252 functional volumes). Six different sequences were programmed and displayed by the Galileo system to counterbalance the order of tactile stimulation between the runs. The marmosets were not required to perform any task.

#### 1.3. Scanning

Data acquisition was performed using a 9.4-T, 31-cm horizontal-bore magnet (Varian/Agilent, Yarnton, UK) and Bruker BioSpec Avance III console with the software package Paravision-6 (Bruker BioSpin Corp, Billerica, MA) at the Centre for Functional and Metabolic Mapping at the University of Western Ontario. We used an in-house, custom-built integrated receive coil with five channels (Schaeffer et al., 2019) paired with a custom-built high-performance 15-cm-diameter gradient coil with 400-mT/m maximum gradient strength (xMR, London, CAN, Peterson et al., 2018). We used an in-house quadrature birdcage coil (12-cm inner diameter) for the transmit coil. Prior to each imaging session, the head chamber was filled with a water-based lubricant gel (MUKO SM321N, Canadian Custom Packaging Company, Toronto, Ontario Canada) to reduce the magnetic-susceptibility image artifacts created by the skull-attached chamber (for details, see Schaeffer et al., 2019).

For each animal, a T2-weighted structural image was acquired, to allow an anatomical registration, with the following parameters: repetition time (TR) = 5500ms; echo time (TE) = 53 ms; field of view (FOV)= 51.2 × 51.2 mm; voxel size = 0.133 × 0.133 × 0.5 mm; number of slices = 42 (axial), bandwidth = 50 kHz, GRAPPA acceleration factor = 2. For functional imaging, gradient-echo-based, single-shot echo-planar images (EPIs) covering the whole brain were acquired over multiple sessions (TR = 1500 ms; TE = 15 ms; flip angle = 40°; FOV = 64 × 64 mm; matrix size = 128 × 128; voxel size = 0.5-mm isotropic; number of slices = 42 (axial); bandwidth = 500 kHz; GRAPPA acceleration factor = 2 (anterior-posterior)).

#### 1.4. Analysis

Time series were preprocessed using AFNI (Cox, 1996), FSL (Smith et al., 2004), SPM12 (Wellcome Department of Cognitive Neurology, London, United Kingdom) and ANTS software (Advanced Normalization Tools, Avants et al., 2011). For spatial preprocessing, functional volumes were first reoriented, realigned (to correct and estimate motion parameters) and a mean functional image was created for each session using SPM12. The images were co-registered with the T2-weighted (T2w) structural image (manually skull-stripped) of each individual monkey using the FMRIB’s linear registration tool (FLIRT) of FSL. Functional images were then non-linearly registered to the NIH marmoset brain atlas (Liu et al., 2018) using ANTs. To reduce noise, images were smoothed by a 1.5-mm full-width at half-maximum Gaussian kernel using SPM12. Bold-oxygen-level-dependent (BOLD) response was estimated using a general linear model (GLM) with SPM12 using the canonical hemodynamic response.

Based on the quality of the images (no artifacts), a total of 16 runs were selected for each monkey. Fixed-effect analyses were performed for each condition at both individual and group level (64 runs in total). In all analyses, six motions regressors (three rotations and three translations) were included as covariates of no interest to remove brain motion artifacts. Thanks to the head-fixation system, little to no motion was observed during imaging sessions (see, Schaeffer et al., 2019). To define the face, arm and leg tactile cortical networks, we contrasted the brain activation obtained by face, arm and leg stimulation, respectively, to those obtained by the no-stimuli condition (Figs 2, 3).

**Figure 2:**
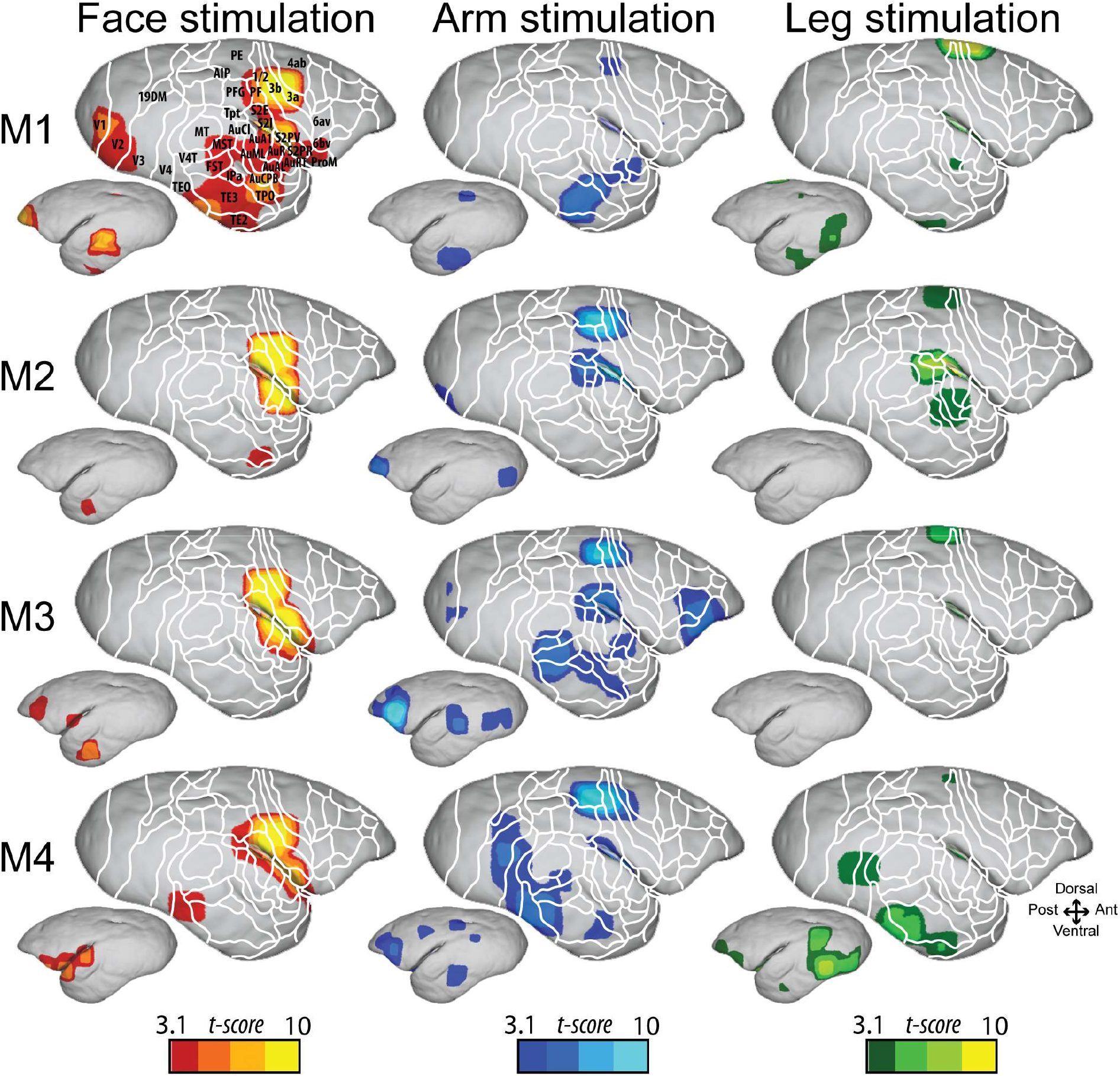
Whole brain mapping of somatosensory responses to face (red), arm (blue) and leg (green) stimulations in each individual marmoset. Each contrast is performed with a level of significance set at p < 0.001 uncorrected level, (t-score > 3.1) Activations are shown on the fiducial brain surface of the NIH marmoset brain atlas for the right contralateral (large brain panel) and left ipsilateral (small brain panel) hemispheres. 1/2, areas 1 and 2 of cortex; 3a, area 3a of somatosensory cortex; 3b, area 3b of somatosensory cortex; 4ab, area 4 of cortex parts a and b; 6av, area 6 of cortex ventral part a; 6bv, area 6 of cortex ventral part b; 19DM, cortex dorsointermediate part; AIP, anterior intraparietal area; AuA1, auditory primary area; AuAL; auditory anterolateral area; AuCl, auditory caudaolateral area; AuCPB, auditory caudal parabelt area; AuML, auditory middle lateral area; AuRT, auditory rostrotemporal area; AuR, auditory rostral area; FST, fundus of the superior temporal sulcus; IPa, intraparietal sulcus associated area; MST, medial superior temporal areas; MT, middle temporal area; PE, parietal area PE; PF, parietal area PF; PFG, parietal area PFG; ProM, proisocortical motor region; S2E, somatosensory area II, external part; S2I, somatosensory area II, internal part; S2PR, somatosensory area II, parietal rostral part; S2PV, somatosensory area II, parietal ventral part; TE2, temporal area TE2; TE3, temporal area TE3; TEO, temporal area TE, occipital part; TPO, temporo-parieto-occipital association area; Tpt, temporoparietal transitional area; V1, visual area 1; V2, visual area 2; V3, visual area 3; V4, visual area 4; V4T, visual area 4 transitional part.

**Figure 3:**
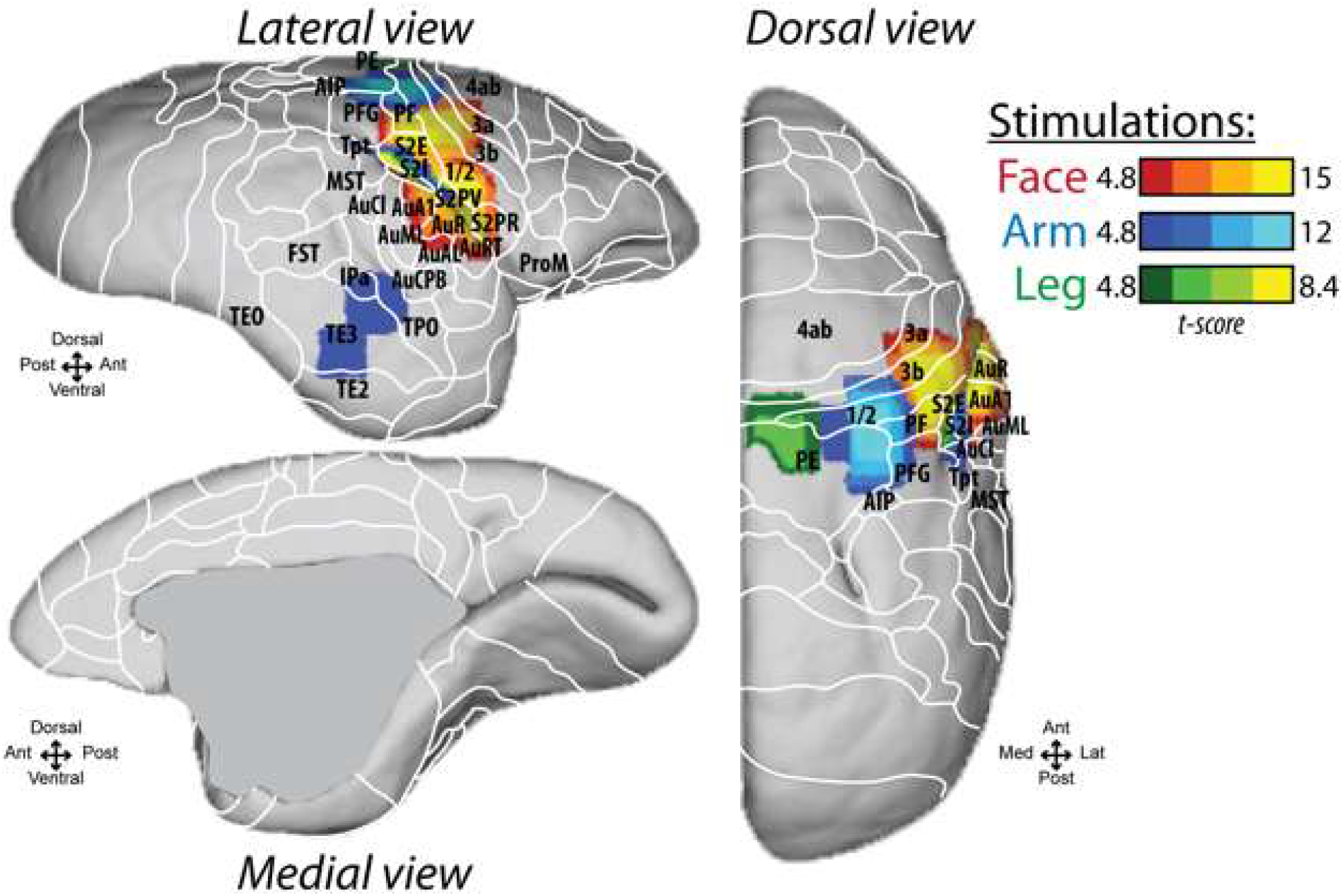
Whole brain mapping of somatosensory responses to face (red), arm (blue) and leg (green) stimulations at the group data set level. Activation are shown on the right fiducial brain surface of the NIH marmoset brain atlas with a level of significance set at p < 0.05 corrected for multiple comparisons (FWE, t-score > 4.8). 23b, cingulate area 23b; 23c, cingulate area 23c; 31, cingulate area 31. All else as in Fig. 2.

#### 1.5. Volumes-of-interest

We performed volumes-of-interest (VOIs) analyses using SPM12, Matlab^®^ (R2018) and the MarsBar toolbox (http://marsbar.sourceforge.net; Brett et al., 2002). VOIs were defined on the group dataset, using face, arm and leg brain activation contrasts obtained at a level of significance set at p < 0.05 corrected for multiple comparisons (Family-wise error: FWE). For each contrast, two VOIs were created by a sphere with a 2mm diameter centered on the local maximum t-score, in SI and SII areas. For each of these six VOIs, the percentage of signal change (PSC) was extracted at their associated coordinates for each run and condition for the group dataset and each individual monkey (Figure 4). The significance of these PSCs across all runs was assessed using paired t-tests for each VOI, after the data were tested for normality (Kolmogorov–Smirnov test, p<0.001 for all VOIs).

**Figure 4:**
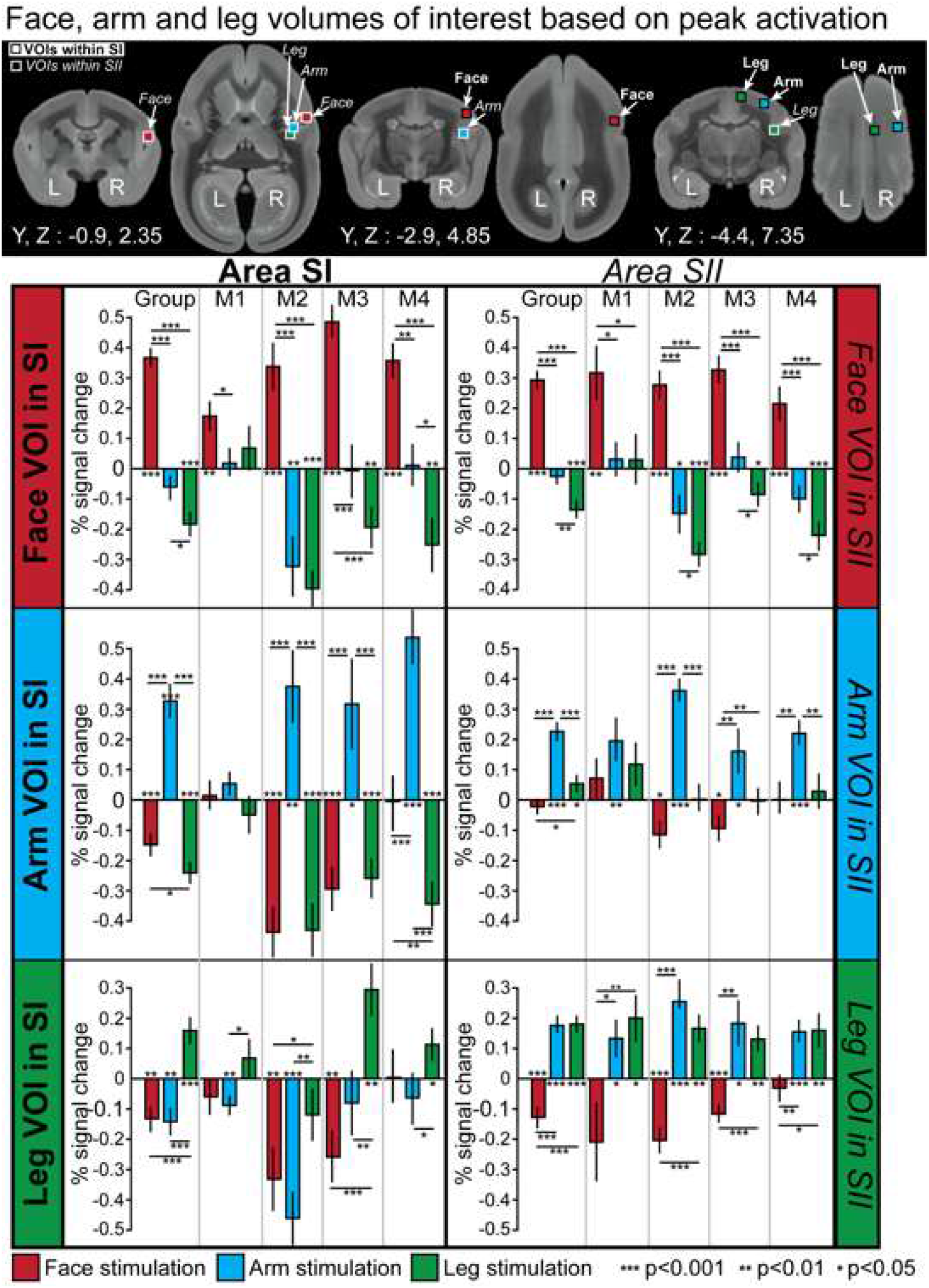
Analysis of percent signal change (PSC) with respect to the baseline (no-stimuli condition) for face, arm and leg stimulation. Upper panel, position of the volumes of interest displayed in coronal and axial views. Squares with black contours correspond to the VOIs defined within the region of SI related to face (red), arm (blue) and leg (green) activation. Squares with white contours correspond to the VOIs defined within the region of SII related to face (red), arm (blue) and leg (green) activation. Lower panel, percent signal change for face, arm and leg VOIs within area SI and SII during face (red), arm (blue) and leg (green) stimulation (paired t-test, *: p<0.05, **: p< 0.01, ***p<0.001). L: left hemisphere; R: right hemisphere.

### 2. Somatosensory system through resting-state-fMRI study

#### 2.1. Subjects and scanning

Five common marmosets (same monkeys as above plus an additional male subject, 36 months, 380 g) were used in RS-fMRI study. Functional images were acquired with 6-22 functional runs (at 400 or 600 volumes each) for each animal in the awake condition, using gradient-echo based single-shot echo-planar imaging sequence with the following parameters: TR = 1500 ms, TE = 15 ms, flip angle = 40°, field of view (FOV) = 64 × 64 mm, matrix size 128 × 128, voxel size 0.5 mm isotropic, slices = 42, bandwidth = 500 kHz, generalized autocalibrating parallel acquisition (GRAPPA) acceleration factor (anterior-posterior) = 2. Total scan time for all resting-state fMRI scans was ~15h.

#### 2.2. Analysis

Data was preprocessed using FSL software (Smith et al., 2004). Functional volumes were first converted, reoriented and brain masks were created using FSL tools and the NIH marmoset T2w brain template (Liu et al., 2018). For each animal, the brain-skull boundary was first roughly identified from individual T2w using the brain extraction tool (BET) with the following options; radius of 25-40 and fractional intensity threshold of 0.3 (Smith, 2002). Then, the NIH T2w brain template was linearly and non-linearly registered to the individual brain image using FMRIB’s linear registration tool (FLIRT) and FMRIB’s nonlinear registration tool (FNIRT) to more accurately create the brain mask. After that, the brain was extracted using the brain mask. RS-fMRI images were corrected for motion using FLIRT. Principal component analysis (PCA) was applied to remove the unstructured noise from the RS-MRI time courses, followed by independent component analysis (ICA) with the decomposition number of 200 using Multivariate Exploratory Linear Optimized Decomposition into the Independent Components (MELODIC) module of the FSL software package. Obtained components were classified as signal or noise (such as eye movement, CSF pulsation, heart rate, and respiratory artifacts) based on the criteria as shown by a previous report (Griffanti et al., 2017), and noise components were regressed out from the RS-fMRI time course using FSL tool (fsl_regfilt). All RS-fMRI images were finally normalized to the NIH template using RS-fMRI-to-T2w and T2w-to-template transformation matrices obtained by FLIRT and FNIRT, followed by spatial smoothing by Gaussian kernel with the full width of half maximum value of 1.0 mm.

Seed-based functional connectivity maps were obtained for each scan by calculating the correlation coefficients (z-transformed) between the time courses in each VOI defined by the task-based MRI analysis (see above “Volumes of interests”) and the time courses in each thalamic voxel (Fig. 6) or in each medial frontal voxel (Fig. 7) using FSL’s FEAT. The functional connectivity maps in the thalamic areas or medial frontal areas, were then averaged across scans and normalized to be maximum z-values equal to 1.

#### 2.3. Delimitation of thalamic nuclei

To create volume of interests (VOIs) in thalamic nuclei, we acquired ex-vivo MRI as it allowed for longer scanning time at a much higher resolution (0.1 mm isotropic). To prepare for ex-vivo MRI, one marmoset (female, 16 months, 306 g) was euthanized through transcardial perfusion and its brain was extracted at the end of the procedure. Anesthesia was initially induced with 30 mg/kg of ketamine and maintained with 4% isoflurane in 1.5-2% oxygen. The animal was then transcardially perfused with 0.9% sodium chloride solution, followed by 10% formaldehyde buffered solution (formalin). The brain was then extracted and stored in 10% buffered formalin for over a week. For ex-vivo imaging, the formalin-fixed marmoset brain was put in a lubricant (Christo-lube; Lubrication technology Inc., Franklin Furnace, OH) to avoid distortion artifact, and three-dimensional multi-echo spin-echo images were acquired with following parameters: TR = 200 ms, TE = 3.5, 8.5, 13.5, 18.5, 23.5 ms, FOV = 33 × 28.8 × 36 mm, matrix size = 330 × 288 × 360, voxel size = 0.1 mm isotropic resolution, averages=4. The average image across different TE images was calculated to increase the signal-to-noise ratio (SNR). This image was also normalized to the NIH template using FLIRT and FNIRT as described above, then, we manually created the subthalamic VOIs (anterior nucleus, laterodorsal nucleus (LD), mediodorsal nucleus(MD), ventral anterior nucleus (VA), ventral lateral nucleus(VL), ventral posterior nucleus (VP) and pulvinar nucleus) with reference to the Paxinos atlas (Paxinos et al., 2011).

When coordinates are provided in this manuscript, they are presented with respect to the anterior commissure. Results are displayed on coronal sections, parasagittal sections or fiducial maps obtained with Caret (Van Essen et al., 2001; http://www.nitrc.org/projects/caret/) using the NIH marmoset brain template (Liu et al., 2018). The labelling refers to the histology-based atlas of Paxinos and colleagues (2011) for the cortical and subcortical regions (Figs. 2, 3, 5 and 6).

**Figure 5:**
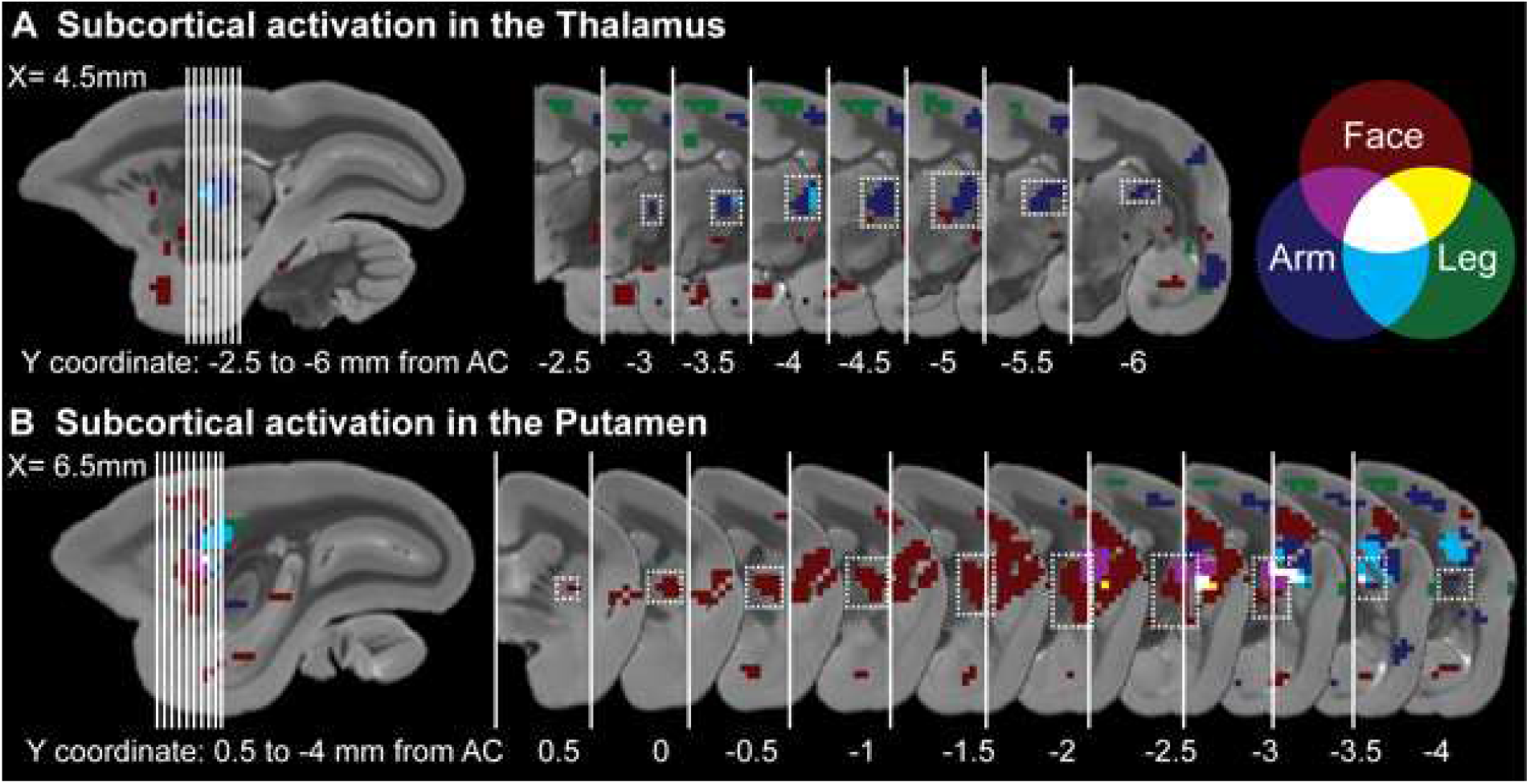
Subcortical activation in the thalamus (A) and putamen (B) at the group level. Face, arm and leg activation are shown in red, blue and green respectively in coronal and sagittal slices (t-score>3.1 for face, arm, leg in putamen and arm in thalamus, t-score > 2.0 for face and leg in thalamus). Overlapping activations are shown in purple for face/arm, in light blue for leg/arm, in yellow for face/leg and in white for face/arm/leg overlapping activation. The white dotted area indicated the activation within the thalamus (A) and putamen (B). Y coordinates are calculated with respect to the anterior commissure (AC).

**Figure 6:**
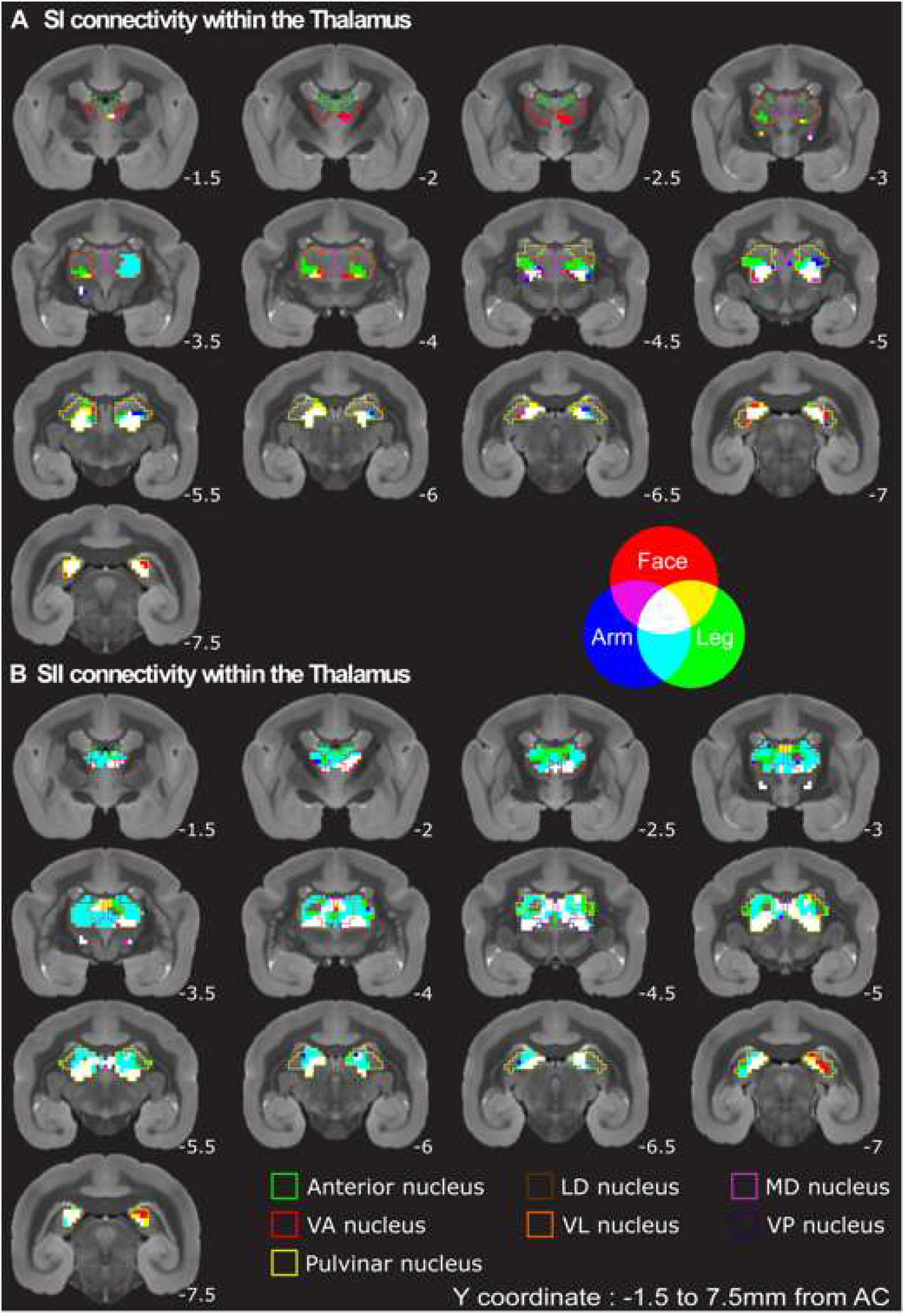
Functional connectivity maps of primary (SI) and secondary somatosensory (SII) areas with thalamic nuclei. Maps are displayed on coronal slices. Face, arm and leg connections are shown in red, blue and green. Overlapping connections are shown in purple for face/arm, in light blue for leg/arm, in yellow for face/leg and in white for face/arm/leg. Contours color matching correspond to green for the anterior nucleus, brown for lateral dorsal (LD) nucleus, pink for medial dorsal (MD) nucleus, red for ventral anterior (VA) nucleus, orange for ventral lateral (VL) nucleus, purple for ventral posterior(VP) nucleus and yellow for pulvinar nucleus. Y coordinates are calculated with respect to the anterior commissure (AC).

## Results

In the following, we describe the functional cortical and subcortical networks activated by tactile stimulation of the face, arm, or leg.

Both face, arm and leg stimulations elicited brain activation in the four marmosets in the contralateral (right) hemisphere of the stimulation (Fig. 2, large brain panel). Tactile activation was found in two main somatosensory regions: SI and SII. However, the position of the activation within the areas differs between the face, arm and leg conditions. Along SI, comprising the areas 1/2, 3a and 3b, face stimulation activated the lateral lower part, arm stimulation activated the middle part and leg stimulation the most upper and medial part of SI. In the same way, within the lateral sulcus, we observed activation in the middle/rostral part for the face and more caudally for the arm and the leg (Fig. 2). Although the general contralateral somatotopic organization looks similar across the animals, it should be noted that the size of activation, the peak location and strength as well as the defined borders slightly vary across the four animals (Fig. 2, Table 1). The pattern of ipsilateral activation was different between monkeys for the same tactile stimulation (Fig. 2, small brain panel). Most of this activation was observed in the temporal cortex. Only marmosets M3 and M4 exhibited tactile activation within the ipsilateral SII. As ipsilateral activation was not consistent across marmosets showing large variabilities in localization and strength of signal, we report only contralateral (right hemisphere) stimulation-evoked activation in the following analyses (Figs. 3, 4, 5 and 7; Tables 1 and 2).

**Figure 7:**
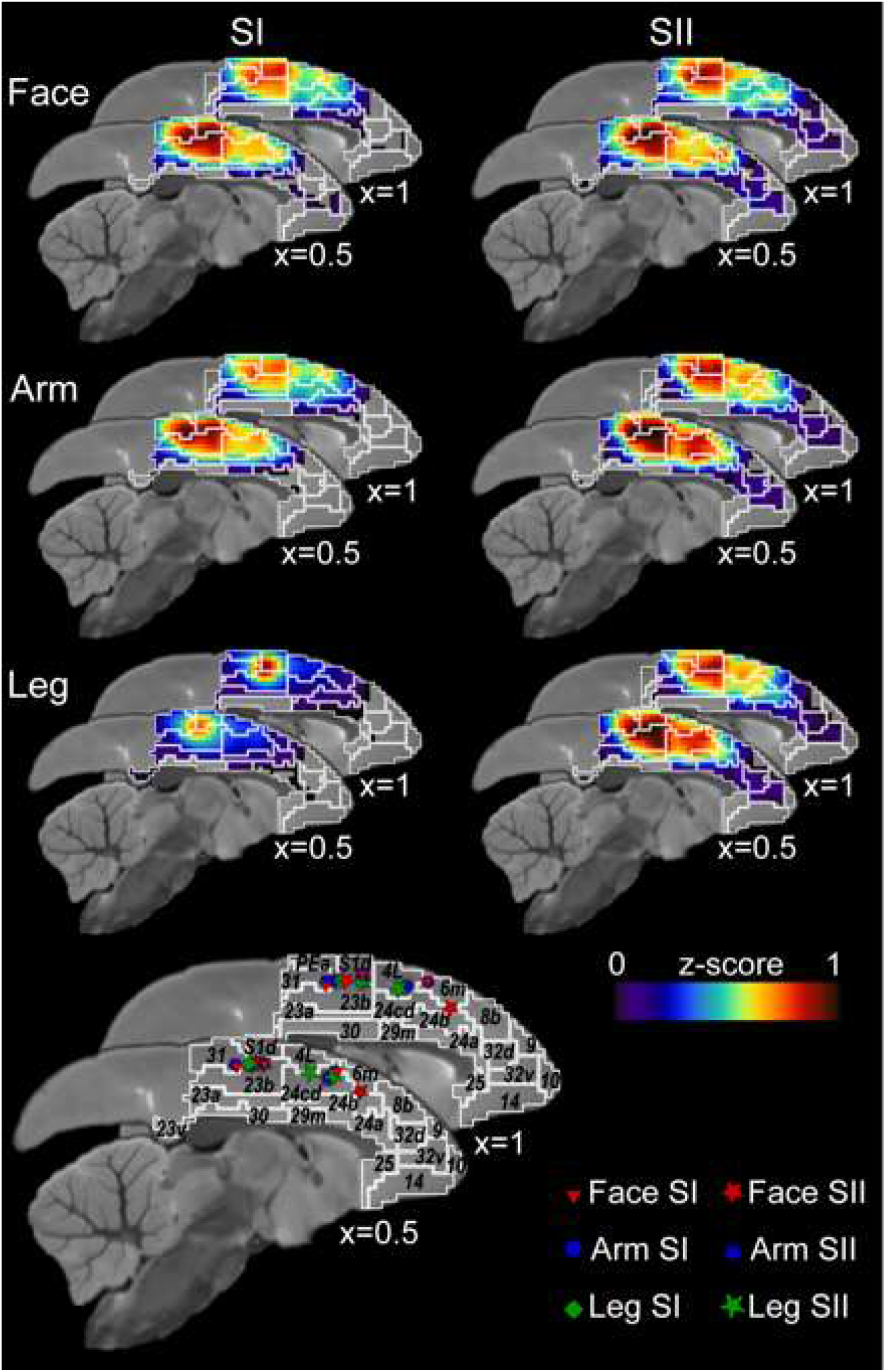
Functional connectivity maps of primary (SI) and secondary (SII) somatosensory areas with medial frontal cortex. Maps show normalized z-scores displayed on parasagittal slices of the right hemisphere. Lower figure shows the maximal peak location for face, arm and leg combined on one brain. White contours show the boundaries of cortical areas. X coordinates are calculated with respect to the anterior commissure (AC). Labelling refers to the connectivity-based parcellation of the NIH marmoset brain atlas (Liu et al., 2018).

**Table 1:** Localization and t-score of the activation peak across the four animals.

| Activation peak | Areas | Animal | Coordinates X, Y, Z<br>in mm | t-score at the peak |
| --- | --- | --- | --- | --- |
| Face | Area SI | M1 | [6.75; -1.40; 5.85] | 10.31 |
|  |  | M2 | [8.25; -1.90; 4.85] | 11.36 |
|  |  | M3 | [7.75; -2.40; 5.35] | 9.67 |
|  |  | M4 | [8.25; -1.90; 4.85] | 9.1 |
|  | Area SII | M1 | [8.75; -1.40; 2.35] | 8.45 |
|  |  | M2 | [9.75; -1.40; 1.85] | 10.16 |
|  |  | M3 | [9.25; -1.40; 2.35] | 11.49 |
|  |  | M4 | [8.75; -0.40; 1.35] | 7.73 |
| Arm | Area SI | M1 | [4.75; -2.90; 7.35] | 3.55 |
|  |  | M2 | [5.75; -4.40; 6.35] | 8.81 |
|  |  | M3 | [4.25; -3.90; 6.85] | 8.46 |
|  |  | M4 | [5.25; -4.40; 6.85] | 10.09 |
|  | Area SII | M1 | [7.25; -2.90; 2.35] | 3.79 |
|  |  | M2 | [7.75; -3.40; 2.35] | 10.52 |
|  |  | M3 | [7.75; -3.90; 3.85] | 5.87 |
|  |  | M4 | [7.25; -2.40; 2.35] | 6.23 |
| Leg | Area SI | M1 | [0.75; -2.90; 8.85] | 10.91 |
|  |  | M2 | [2.75; -4.90; 7.85] | 4.13 |
|  |  | M3 | [1.75; -4.40; 7.85] | 5.43 |
|  |  | M4 | [2.75; -3.90; 7.85] | 3.27 |
|  | Area SII | M1 | [6.25; -4.40; 3.85] | 6.11 |
|  |  | M2 | [7.75; -4.90; 2.85] | 9.76 |
|  |  | M3 | [7.25; -3.90; 3.35] | 4.5 |
|  |  | M4 | [6.75; -3.90; 2.85] | 4.65 |

**Table 2:** Coordinates and t-score of the volumes of interest selected for the PSC analysis (based on the NIH marmoset brain atlas and the group level analysis).

| Activation peak | Areas | Coordinates X, Y, Z<br>in mm | t-score at the peak |
| --- | --- | --- | --- |
| Face | Area SI | [7.75; -2.40; 5.35] | 13.13 |
|  | Area SII | [9.25; -1.40; 1.85] | 14.84 |
| Arm | Area SI | [5.25; -3.90; 6.85] | 10.32 |
|  | Area SII | [7.25; -2.90; 2.35] | 9.97 |
| Leg | Area SI | [1.75; -4.40; 7.85] | 5.71 |
|  | Area SII | [6.75; -3.90; 2.85] | 9.13 |

Figure 3 shows the activation maps at the group level (64 runs) obtained by the same analysis. It highlights the medio-lateral progression from the leg to the face and the rostro-caudal progression from the face to the leg for the body tactile representation in SI and SII, respectively, observed in both individual (Fig. 2) and group level analyses (Fig. 3).

### Face stimulation mapping

The air-puff delivery to the face led to contralateral activation in the somatosensory area SI with areas 1/2, 3a, 3b; somatosensory area SII external part (S2E), internal part (S2I), parietal rostral area (S2PR), parietal ventral area (S2PV); parietal areas PF and PFG; proisocortical motor region (ProM), dysgranular (DI) and granular (GI) insular cortex; retroinsular area (ReI); temporal area TE3. We also observed activation within the contralateral auditory cortex with auditory primary area (AuA1), anterolateral area (AuAL), caudomedial area (AuCM), rostral area (AuR), rostrotemporal area (AuRT), rostromedial area (AuRM), rostrotemporal medial area (AuRMT). Subcortical activation were found in the contralateral hippocampus, amygdala, thalamus, caudate and putamen. Some weak activation were also found in the hemisphere ipsilateral to stimulation in the caudate and putamen.

### Arm stimulation mapping

Tactile stimulation performed to the arm led to activation in the somatosensory area SI with areas 1/2, 3a, 3b; somatosensory area SII external part (S2E), internal part (S2I), parietal ventral area (S2PV); parietal areas PE, PF and PFG; granular insular cortex (GI); retroinsular area (ReI); temporal areas TE2, TE3, TE occipital part (TEO); temporo-parieto-occipital association area (TPO); temporoparietal transitional area (Tpt); fundus of superior temporal sulcus area (FST); medial superior temporal area (MST); intraparietal sulcus associated area (IPa); auditory caudomedial area (AuCM). Subcortical activation was found in the thalamus, caudate and putamen. Some weak activation was also found in the hemisphere ipsilateral to stimulation in the hippocampus, caudate and putamen.

### Leg stimulation mapping

The leg stimulation activated the somatosensory area SI with areas 1/2, 3a, 3b; somatosensory area SII internal part (S2I); parietal area PE; granular insular cortex (GI); retroinsular area (ReI); temporoparietal transitional area (Tpt); auditory caudomedial area (AuCM), middle lateral area (AuML), caudal parabelt area (AuCPB), areas 4ab, 31, 23b, 23c. Subcortical activations were found in the hippocampus, thalamus, caudate and putamen.

### Volumes of interest

Based on the whole brain mapping of somatosensory responses performed at the group level, six volumes of interest (VOI) were defined in SI and SII, corresponding to face, arm and leg activation local maxima in these areas. VOIs were created by a 2mm of diameter sphere centered on the activation peak (Table 2). The PSC was extracted for each region for each monkey individually and at the group level (Fig. 4).

The activation peak coordinates (Table 2, Fig. 5, upper panel) validated the medio-lateral and rostro-caudal progressions observed for the body tactile representation in SI and SII in the contrast maps. The PSC analysis (Fig. 5, lower panel) showed the face VOIs within area SI and area SII exhibited significantly higher PSC during the face stimulation compared to the arm and leg stimulation. The same pattern was observed for the arm VOIs within area SI and area SII but, this case, in favor of the arm stimulation. The leg VOIs showed more variability, PSC in area SI were higher during leg stimulation but in area SII, PSC were high during both arm and leg stimulation and did not show any difference between these two conditions. However, the PSC during the face stimulation were significantly negative.

Overall, in area SI, PSCs were high during the related body part stimulation and negative for the other body part stimulation. In area SII, PSCs were high for the face during face stimulation and for the arm during arm stimulation but lower compared to SI. Interestingly, the PSC extracted at the group level reflected quite well the PSC extracted at the individual level even though some variability can be observed across animals.

### Subcortical activation

Subcortical activation in the thalamus and putamen showed some somatotopy of the different body part stimulation (Fig. 5). In the thalamus (Fig. 5A), arm stimulation led to the strongest activation (dark blue, t-score > 3.1) compared to the face and leg stimulation (red and green, respectively, t-score >2). We observed arm and leg activation in the lateral part of the thalamus and face activation in a more medial and posterior part of the thalamus. The leg activation is fully included in the area activated by arm activation (arm/leg overlapping activation are in light blue color). Referring to the histology-based atlas of Paxinos (2011), the face activation area corresponds to the ventral posteromedial thalamic nucleus (VPM) and the arm and leg activation to the ventral posterolateral thalamic nucleus (VPL).

In the putamen (Fig. 5B), face stimulation led to the strongest activation compared to the arm and leg stimulation (red, blue and green, respectively, t-score >3.1). We observed face activation more rostral than arm and leg activation. The leg activation was more lateral and posterior than the arm and face activation. There was a strong overlap between the three activations (white), face and arm activation (purple), face and leg activation (yellow) and arm and leg activation (light blue). This overlap suggests that even though we observed a ventrolateral to dorsomedial progression from the face to the leg representation, activation were not exclusive to each body part representation.

### Functional connectivity

Next, we investigated the functional connectivity of six different VOIs, defined previously by the task-related activation, within the thalamic region (Fig. 6) and the medial frontal cortices using resting-state fMRI data (Fig. 7).

For the thalamic connectivity with VOIs defined in SI (Fig. 6A), the face VOI was strongly connected with the ventromedial part, the arm VOI with the lateral part and the leg VOI with the most dorsolateral part of the thalamus. Face, arm and leg VOIs showed a strong overlap in their connectivity map within the pulvinar and ventral posterior (VP) nucleus. The ventral anterior (VA) nucleus had mostly functional connectivity with the face VOI, and the ventral lateral (VL) nucleus with arm and leg VOIs.

For the thalamic connectivity with VOIs defined in SII (Fig. 6B), the connectivity maps were larger than for SI VOIs. A strong overlap was observed in lower parts of the VA and VL nuclei, in the VP nucleus and medioventral part of the pulvinar. Arm and leg VOIs did not exhibit specific functional connectivity related to the body part as their connectivity maps mainly overlapped. Face VOI showed specific connections with the posterior part of the pulvinar.

For the medial frontal cortex (Fig. 7), VOIs defined in SI showed strong connectivity with the posterior regions (areas 23b and 31). The face and arm VOIs exhibited strong connectivity with midcingulate areas (area 24b, c and d). VOIs defined in SII showed strong connectivity with both posterior and midcingulate cortex for face, arm and leg. Interestingly, for midcingulate areas, the hot spot of connectivity was more anterior for the face compared to the arm and leg, suggesting a body representation mapping within the midcingulate cortex. In addition, face VOIs were functionally connected with the medial part of area 6 whereas arm and leg VOIs showed connections with the lateral part of motor area 4. Overall, face VOIs exhibited stronger functional connectivity profile than arm and leg VOIs. VOIs defined in SII showed higher connectivity maps than for VOIs defined in SI.

## Discussion

In this study, we identified the somatotopic representation of tactile stimulation in awake marmosets by using ultrahigh-field 9.4T fMRI. By stimulating the face, arm or leg of marmoset, we showed strong contralateral functional activation related to each body part and defined topographic body representations in somatosensory areas SI, SII. These activation areas were generally consistent with previous findings obtained by invasive methods. We also examined the subcortical activation patterns associated with tactile stimulation and highlighted that the thalamus and, for the first time, the putamen exhibited somatotopic patterns in marmosets. Using RS-fMRI, we also revealed how somatosensory areas highly responsive to tactile stimulation are functionally connected to other cortical and subcortical regions. These connections were consistent with previous findings in humans and macaque monkeys, indicating that the organization of the somatosensory system is conserved across primates.

### Somatotopy in the primary and secondary somatosensory regions

The primary somatosensory cortex (SI) encompasses areas 3a, 3b, 1 and 2 in primates (Merzenich et al., 1978; Sur et al., 1982; Kaas, 1983, 1987, 1989; Krubitzer, 1996). We observed face, arm and leg activation in all of these areas but in different locations: face activation were in the lateral lower part, the arm activation in the middle part and leg activation in the most medial and dorsal part of SI. This is in accordance with previous recordings and anatomical studies showing the existence of a body surface representation in SI from the foot in the most medial part to the face and mouth representations in the most lateral part, other body part representations such as leg, trunk, arm, hand, digits successively followed a mediolateral sequence (Garraghty et al., 1990; Krubitzer and Kaas, 1990, 1992; Huffman and Krubitzer, 2001a, 2001b). This pattern has been identified in area 3b (Krubitzer and Kaas, 1988, 1990, 1992; Garraghty et al., 1990), in area 3a, located rostrally (Krubitzer and Kaas, 1988; Garraghty et al., 1990; Huffman and Krubitzer, 2001a, 2001b) and in area 1/2 located caudally to 3b but reversely mirroring the pattern of 3b (Krubitzer and Kaas, 1988, 1990; Kaas, 2004a).

We observed tactile activation within the lateral sulcus: in the middle/rostral part for the face and more caudally for the arm and the leg representations. This is in accordance with the somatotopic representation of the body surface following a rostrocaudal progression deeper in the sulcus from the face to the tail described by electrophysiology in anesthetized marmosets for SII (Krubitzer and Kaas, 1990; Kaas, 2004b, 2004a) and PV (Krubitzer and Kaas, 1990; Qi et al., 2002).

As our tactile stimulation corresponded to the delivery of air-puffs, we expected to activate mostly cutaneous receptors found in 3b, SII and PV (Krubitzer and Kaas, 1988, 1990; Qi et al., 2002). Interestingly, we also observed large activation in 3a and 1/2 which were slightly or not receptive to skin stimulation in previous studies but found to be densely interconnected with 3b, SII and PV (Huffman and Krubitzer, 2001a). As fMRI reflects both neuronal activities and modulation (Logothetis et al., 2001; Logothetis and Pfeuffer, 2004; Magri et al., 2012), these activation can reflect the connectivity network.

### Overlapping between representation

In SI we observed positive PSC only for the dedicated representation whereas in SII, we observed positive PSCs for both arm and leg stimulation within the leg representation. The overlap between representations suggests that SII did not have a specific body part representation. Some studies in humans have shown a representational overlap between digits representations linked to a common use of these digits (Ejaz et al., 2015) both in active and passives tasks (Berlot et al., 2018; Sanders et al., 2019), but much stronger in SII than SI (Simões et al., 2001; Sanchez Panchuelo et al., 2018). Our results suggest this overlapping representation can be extended to other body parts in marmosets.

### Functional connectivity with the medial frontal cortex

Our functional connectivity maps between SI/SII and the medial frontal cortex highlight two main subfields in body representation, one in the posterior cingulate cortex and one more rostrally. Interestingly, a somatomotor map for face, upper and lower limb has been identified in rostral cingulate motor area (Misra and Coombes, 2015) and more posteriorly both in humans and monkeys (Wang et al., 2004; Morecraft and Tanji, 2009; Amiez and Petrides, 2014). Our results suggest that a similar organization is present in marmosets. In addition, these subfields have been proposed to play a role in embodied representation of feedback in humans and macaques, as multiple activations related to eye movement, blink, juice reward or tactile stimulation overlap in two separate subfields (Amiez et al., 2013; Procyk et al., 2014; Cléry et al., 2017a).

### Somatotopy in subcortical areas: thalamus and putamen

Activation in the lateral part (VPL) and in the medial and posterior part of the thalamus (VPM) were observed after we stimulated the arm/leg and face, respectively. This somatotopic organization followed a mediolateral sequence from face to lower limbs previously described in marmosets ventroposterior nucleus VP (Krubitzer and Kaas, 1992; Wilson et al., 1999) and in other species including non-human primates and humans (for reviews, see Nambu, 2011; Chien et al., 2017).

Both SI and SII were connected to the thalamus but exhibited a different pattern of functional connectivity. As for the activation maps, we observed a ventromedial-dorsolateral organization from face to leg. However, strong overlaps in SI connectivity maps were found in pulvinar and VP suggesting that parts of these areas process tactile information regardless of the stimulated body part. VP has been shown to be strongly connected to 3a and 3b (Brysch et al., 1990; Krubitzer and Kaas, 1992; Huffman and Krubitzer, 2001b; Hori et al., 2020b). SII connectivity maps showed mainly overlap between body representations. However, VPi, anatomically connected to 3a, SII and PV, was found to show a body representation paralleling that in VP (Huffman and Krubitzer, 2001b; Qi et al., 2002). This difference between our results and anatomical studies may be due to our current MRI resolution leading to weak thalamic signals and precluding deeper analyses.

This is the first study showing a somatotopic organization within the putamen in marmosets. This ventrolateral to dorsomedial progression from the face to the tail representation has been shown in Old Word macaque monkeys and humans (Romanelli et al., 2005; Nambu, 2011; Zeharia et al., 2015) and seems to be present in New World marmosets, too. A somatotopic organization linked to the different projections received by the putamen has been described (for reviews, see Romanelli et al., 2005; Nambu, 2011). The primary motor area M1, supplementary motor area SMA, the motor thalamus and the two cingulate motor areas project to the putamen in independent regions and/or convergence areas while following a somatotopic ventrolateral-dorsomedial progression (Romanelli et al., 2005; Nambu, 2011). Our results suggest that the putamen plays a key role in the sensorimotor processing in the marmoset brain, even during passive tactile stimulation. Further studies on how these areas project to the marmoset’s putamen need to be investigated.

### Limitations

In the current task, we stimulated only passively the left side due to set-up and acquisition constraints (space, time…) however we expect that the stimulation of the right side will lead to a similar body representation pattern in the contralateral (left) hemisphere (Liu et al., 2013). As active and passive stimulations can elicit the same pattern of activation in the human contralateral side (Berlot et al., 2018; Sanders et al., 2019), it would be interesting to explore if it applies to marmosets. In addition, a recent study in human subjects has shown that ipsilateral fingers representations were mainly driven by an active movement rather than a passive stimulation (Berlot et al., 2018). With our passive task, we observed weak activation in the ipsilateral SII of marmosets 3 and 4 and mainly deactivation within the ipsilateral SI for all marmosets. This ipsilateral SI suppression in favor of the contralateral side was mainly studied for fingers and hands representation in humans (Staines et al., 2002; Hlushchuk and Hari, 2006; Del Vecchio et al., 2019; Lamp et al., 2019) but seems to be generalized to the whole body in marmosets. Further studies testing active and passive tactile tasks using left and right stimulations (alone or simultaneously) should be performed.

Solenoid valves opening from the tactile stimulation system might have been perceived by the marmosets as it was a punctual noise compared to the constant MR scanner sound during the run and could explain why we observed functional activation in auditory regions. As the tube was close to the face, it might have increased the probability of hearing that sound by marmosets during face stimulation.

### Perspectives

This non-invasive method will be useful to map target regions in individual marmosets for guiding future invasive recordings and neural manipulation techniques (e.g. electrophysiological recordings, calcium imaging, optogenetics). This whole brain mapping can be performed for a more detailed body representation and combined to other sensory modalities (visual, auditory) to explore multisensory integration process (Cléry et al., 2015, 2017b, 2020), or to functional tasks to explore motor-related function such as grasping (Tia et al., 2017).

## Acknowledgements

Support was provided by the Canadian Institutes of Health Research (FRN 148365), and the Canada First Research Excellence Fund to BrainsCAN, and Canada Research Chair. We thank Miranda Bellyou, Cheryl Vander Tuin, and Whitney Froese for animal preparation and care and Dr. Alex Li for scanning assistance. We also thank Ruth Lanius for lending us the Galileo somatosensory stimulus system.

## Notes

**Conflict of interest:** The authors declare no conflict of interest.

### Competing Interest Statement

The authors have declared no competing interest.

## References

Amiez C, Neveu R, Warrot D, Petrides M, Knoblauch K, Procyk E (2013) The Location of Feedback-Related Activity in the Midcingulate Cortex Is Predicted by Local Morphology. J Neurosci 33:2217–2228.

Amiez C, Petrides M (2014) Neuroimaging Evidence of the Anatomo-Functional Organization of the Human Cingulate Motor Areas. Cereb Cortex 24:563–578.

Avants BB, Tustison NJ, Song G, Cook PA, Klein A, Gee JC (2011) A reproducible evaluation of ANTs similarity metric performance in brain image registration. Neuroimage 54:2033–2044.

Berlot E, Prichard G, O’Reilly J, Ejaz N, Diedrichsen J (2018) Ipsilateral finger representations in the sensorimotor cortex are driven by active movement processes, not passive sensory input. Journal of Neurophysiology 121:418–426.

Brett M, Anton J-L, Valabregue R, Poline J-B (2002) Region of interest analysis using the MarsBar toolbox for SPM 99. Presented at the 8th International Conference on Functional Mapping of the Human Brain (June 2-6, 2002, Sendai, Japan), 16:S497.

Brysch W, Brysch I, Creutzfeldt OD, Schlingensiepen R, Schlingensiepen KH (1990) The topology of the thalamo-cortical projections in the marmoset monkey (Callithrix jacchus). Exp Brain Res 81:1–17.

Burkart JM, Finkenwirth C (2015) Marmosets as model species in neuroscience and evolutionary anthropology. Neuroscience Research 93:8–19.

Carlson M, Huerta MF, Cusick CG, Kaas JH (1986) Studies on the evolution of multiple somatosensory representations in primates: The organization of anterior parietal cortex in the new world callitrichid, Saguinus. Journal of Comparative Neurology 246:409–426.

Chien JH, Korzeniewska A, Colloca L, Campbell C, Dougherty P, Lenz F (2017) Human Thalamic Somatosensory Nucleus (Ventral Caudal, Vc) as a Locus for Stimulation by INPUTS from Tactile, Noxious and Thermal Sensors on an Active Prosthesis. Sensors (Basel) 17

Cléry J, Amiez C, Guipponi O, Wardak C, Procyk E, Ben Hamed S (2017a) Reward activations and face fields in monkey cingulate motor areas. Journal of Neurophysiology 119:1037–1044.

Cléry J, Guipponi O, Odouard S, Pinède S, Wardak C, Hamed SB (2017b) The Prediction of Impact of a Looming Stimulus onto the Body Is Subserved by Multisensory Integration Mechanisms. J Neurosci 37:10656–10670.

Cléry J, Guipponi O, Odouard S, Wardak C, Ben Hamed S (2015) Impact prediction by looming visual stimuli enhances tactile detection. J Neurosci 35:4179–4189.

Cléry JC, Schaeffer DJ, Hori Y, Gilbert KM, Hayrynen LK, Gati JS, Menon RS, Everling S (2020) Looming and receding visual networks in awake marmosets investigated with fMRI. NeuroImage 215:116815.

Cox RW (1996) AFNI: software for analysis and visualization of functional magnetic resonance neuroimages. Comput Biomed Res 29:162–173.

Del Vecchio M, Caruana F, Sartori I, Pelliccia V, Lo Russo G, Rizzolatti G, Avanzini P (2019) Ipsilateral somatosensory responses in humans: the tonic activity of SII and posterior insular cortex. Brain Struct Funct 224:9–18.

Ejaz N, Hamada M, Diedrichsen J (2015) Hand use predicts the structure of representations in sensorimotor cortex. Nature Neuroscience 18:1034–1040.

Garraghty PE, Pons TP, Kaas JH (1990) Ablations of areas 3b (SI proper) and 3a of somatosensory cortex in marmosets deactivate the second and parietal ventral somatosensory areas. Somatosens Mot Res 7:125–135.

Griffanti L, Douaud G, Bijsterbosch J, Evangelisti S, Alfaro-Almagro F, Glasser MF, Duff EP, Fitzgibbon S, Westphal R, Carone D, Beckmann CF, Smith SM (2017) Hand classification of fMRI ICA noise components. Neuroimage 154:188–205.

Hirano Y, Yen CC, Liu JV, Mackel JB, Merkle H, Nascimento GC, Stefanovic B, Silva AC (2018) Investigation of the BOLD and CBV fMRI responses to somatosensory stimulation in awake marmosets (Callithrix jacchus). NMR in Biomedicine 31:e3864.

Hlushchuk Y, Hari R (2006) Transient Suppression of Ipsilateral Primary Somatosensory Cortex during Tactile Finger Stimulation. J Neurosci 26:5819–5824.

Hori Y, Schaeffer DJ, Gilbert KM, Hayrynen LK, Cléry JC, Gati JS, Menon RS, Everling S (2020a) Comparison of resting-state functional connectivity in marmosets with tracerbased cellular connectivity. NeuroImage 204:116241.

Hori Y, Schaeffer DJ, Yoshida A, Clery JC, Hayrynen LK, Gati JS, Menon RS, Everling S (2020b) Cortico-subcortical functional connectivity profiles of resting-state networks in marmosets and humans. bioRxiv:2020.07.14.202382.

Huffman KJ, Krubitzer L (2001a) Area 3a: Topographic Organization and Cortical Connections in Marmoset Monkeys. Cereb Cortex 11:849–867.

Huffman KJ, Krubitzer L (2001b) Thalamo-cortical connections of areas 3a and M1 in marmoset monkeys. Journal of Comparative Neurology 435:291–310.

Johnston KD, Barker K, Schaeffer L, Schaeffer D, Everling S (2018) Methods for chair restraint and training of the common marmoset on oculomotor tasks. Journal of Neurophysiology 119:1636–1646.

Kaas JH (1983) What, if anything, is SI? Organization of first somatosensory area of cortex. Physiological Reviews 63:206–231.

Kaas JH (1987) The Organization of Neocortex in Mammals: Implications for Theories of Brain Function. Annual Review of Psychology 38:129–151.

Kaas JH (1989) The evolution of complex sensory systems in mammals. Journal of Experimental Biology 146:165–176.

Kaas JH (2004a) Somatosensory Cortex. In: Encyclopedia of Neuroscience (Squire LR, ed), pp 73–77. Oxford: Academic Press.

Kaas JH (2004b) Evolution of somatosensory and motor cortex in primates. The Anatomical Record Part A: Discoveries in Molecular, Cellular, and Evolutionary Biology 281A:1148–1156.

Kaas JH, Nelson RJ, Sur M, Lin CS, Merzenich MM (1979) Multiple representations of the body within the primary somatosensory cortex of primates. Science 204:521–523.

Krubitzer LA (1996) The organization of lateral somatosensory cortex in primates and other mammals. In: Somesthesis and the Neurobiology of the Somatosensory Cortex (Franzén O, Johansson R, Terenius L, eds), pp 173–185 Advances in Life Sciences. Basel: Birkhäuser.

Krubitzer LA, Kaas JH (1988) Responsiveness and somatotopic organization of anterior parietal field 3b and adjoining cortex in newborn and infant monkeys. Somatosens Mot Res 6:179–205.

Krubitzer LA, Kaas JH (1990) The organization and connections of somatosensory cortex in marmosets. J Neurosci 10:952–974.

Krubitzer LA, Kaas JH (1992) The somatosensory thalamus of monkeys: Cortical connections and a redefinition of nuclei in marmosets. Journal of Comparative Neurology 319:123–140.

Lamp G, Goodin P, Palmer S, Low E, Barutchu A, Carey LM (2019) Activation of Bilateral Secondary Somatosensory Cortex With Right Hand Touch Stimulation: A Meta-Analysis of Functional Neuroimaging Studies. Front Neurol 9

Liu C, Ye FQ, Yen CC-C, Newman JD, Glen D, Leopold DA, Silva AC (2018) A digital 3D atlas of the marmoset brain based on multi-modal MRI. NeuroImage 169:106–116.

Liu JV, Hirano Y, Nascimento GC, Stefanovic B, Leopold DA, Silva AC (2013) fMRI in the awake marmoset: Somatosensory-evoked responses, functional connectivity, and comparison with propofol anesthesia. NeuroImage 78:186–195.

Logothetis NK, Pauls J, Augath M, Trinath T, Oeltermann A (2001) Neurophysiological investigation of the basis of the fMRI signal. Nature 412:150–157.

Logothetis NK, Pfeuffer J (2004) On the nature of the BOLD fMRI contrast mechanism. Magnetic Resonance Imaging 22:1517–1531.

Magri C, Schridde U, Murayama Y, Panzeri S, Logothetis NK (2012) The Amplitude and Timing of the BOLD Signal Reflects the Relationship between Local Field Potential Power at Different Frequencies. J Neurosci 32:1395–1407.

Mansfield K (2003) Marmoset models commonly used in biomedical research. Comp Med 53:383–392.

Merzenich MM, Kaas JH, Sur M, Lin CS (1978) Double representation of the body surface within cytoarchitectonic areas 3b and 1 in “SI” in the owl monkey (Aotus trivirgatus). J Comp Neurol 181:41–73.

Misra G, Coombes SA (2015) Neuroimaging Evidence of Motor Control and Pain Processing in the Human Midcingulate Cortex. Cereb Cortex 25:1906–1919.

Morecraft RJ, Tanji J (2009) Cingulofrontal interactions and the cingulate motor areas. Cingulate neurobiology and disease Oxford University Press, UK:113–144.

Nambu A (2011) Somatotopic Organization of the Primate Basal Ganglia. Front Neuroanat 5

Papoti D, Yen CC-C, Mackel JB, Merkle H, Silva AC (2013) An embedded four-channel receive-only RF coil array for fMRI experiments of the somatosensory pathway in conscious awake marmosets. NMR Biomed 26:1395–1402.

Paxinos G, Watson C, Petrides M, Rosa M, Tokuno H (2011) The Marmoset Brain in Stereotaxic Coordinates, 1 edition. Amsterdam; London: Academic Press.

Peterson J, Chaddock R, Dalrymple B, Van Sas F, Gilbert KM, Klassen LM, Gati JS, Handler WB, Chronik BA (2018) Development of a gradient and shim insert system for marmoset imaging at 9.4 T. In: 26th Annual Meeting of International Society for Magnetic Resonance in Medicine, Paris, June.

Preuss TM (2019) Critique of Pure Marmoset. BBE 93:92–107.

Procyk E, Wilson CRE, Stoll FM, Faraut MCM, Petrides M, Amiez C (2014) Midcingulate Motor Map and Feedback Detection: Converging Data from Humans and Monkeys. Cereb Cortex:bhu213.

Qi H-X, Lyon DC, Kaas JH (2002) Cortical and thalamic connections of the parietal ventral somatosensory area in marmoset monkeys (Callithrix jacchus). Journal of Comparative Neurology 443: 168–182.

Romanelli P, Esposito V, Schaal DW, Heit G (2005) Somatotopy in the basal ganglia: experimental and clinical evidence for segregated sensorimotor channels. Brain Research Reviews 48:112–128.

Sanchez Panchuelo RM, Besle J, Schluppeck D, Humberstone M, Francis S (2018) Somatotopy in the Human Somatosensory System. Front Hum Neurosci 12:235.

Sanders Z-B, Wesselink DB, Dempsey-Jones H, Makin TR (2019) Similar somatotopy for active and passive digit representation in primary somatosensory cortex. bioRxiv: 754648.

Sasaki E et al. (2009) Generation of transgenic non-human primates with germline transmission. Nature 459:523–527.

Schaeffer DJ, Gilbert KM, Hori Y, Gati JS, Menon RS, Everling S (2019) Integrated radiofrequency array and animal holder design for minimizing head motion during awake marmoset functional magnetic resonance imaging. NeuroImage 193:126–138.

Silva AC (2017) Anatomical and functional neuroimaging in awake, behaving marmosets. Developmental Neurobiology 77:373–389.

Silva AC, Liu JV, Hirano Y, Leoni RF, Merkle H, Mackel JB, Zhang XF, Nascimento GC, Stefanovic B (2011) Longitudinal Functional Magnetic Resonance Imaging in Animal Models. In: Magnetic Resonance Neuroimaging: Methods and Protocols (Modo M, Bulte JWM, eds), pp 281–302 Methods in Molecular Biology. Totowa, NJ: Humana Press.

Simões C, Mertens M, Forss N, Jousmäki V, Lütkenhöner B, Hari R (2001) Functional Overlap of Finger Representations in Human SI and SII Cortices. Journal of Neurophysiology 86:1661–1665.

Smith SM (2002) Fast robust automated brain extraction. Human Brain Mapping 17:143–155.

Smith SM, Jenkinson M, Woolrich MW, Beckmann CF, Behrens TEJ, Johansen-Berg H, Bannister PR, De Luca M, Drobnjak I, Flitney DE, Niazy RK, Saunders J, Vickers J, Zhang Y, De Stefano N, Brady JM, Matthews PM (2004) Advances in functional and structural MR image analysis and implementation as FSL. Neuroimage 23 Suppl 1:S208–219.

Staines WR, Graham SJ, Black SE, McIlroy WE (2002) Task-Relevant Modulation of Contralateral and Ipsilateral Primary Somatosensory Cortex and the Role of a Prefrontal-Cortical Sensory Gating System. NeuroImage 15:190–199.

Sur M, Nelson RJ, Kaas JH (1982) Representations of the body surface in cortical areas 3b and 1 of squirrel monkeys: Comparisons with other primates. Journal of Comparative Neurology 211:177–192.

Takahashi DY, Fenley AR, Teramoto Y, Narayanan DZ, Borjon JI, Holmes P, Ghazanfar AA (2015) LANGUAGE DEVELOPMENT. The developmental dynamics of marmoset monkey vocal production. Science 349:734–738.

Tia B, Takemi M, Kosugi A, Castagnola E, Ansaldo A, Nakamura T, Ricci D, Ushiba J, Fadiga L, Iriki A (2017) Cortical control of object-specific grasp relies on adjustments of both activity and effective connectivity: a common marmoset study. The Journal of Physiology 595:7203–7221.

Wang Y, Matsuzaka Y, Shima K, Tanji J (2004) Cingulate cortical cells projecting to monkey frontal eye field and primary motor cortex. Neuroreport 15:1559–1563.

Wilson P, Kitchener PD, Snow PJ (1999) Cutaneous receptive field organization in the ventral posterior nucleus of the thalamus in the common marmoset. J Neurophysiol 82:1865–1875.

Workman KP, Healey B, Carlotto A, Lacreuse A (2019) One-year change in cognitive flexibility and fine motor function in middle-aged male and female marmosets (Callithrix jacchus). American Journal of Primatology 81:e22924.

Zeharia N, Hertz U, Flash T, Amedi A (2015) New Whole-Body Sensory-Motor Gradients Revealed Using Phase-Locked Analysis and Verified Using Multivoxel Pattern Analysis and Functional Connectivity. J Neurosci 35:2845–2859.

